## Supplementary Figures for "The structural coverage of the human proteome before and after AlphaFold"

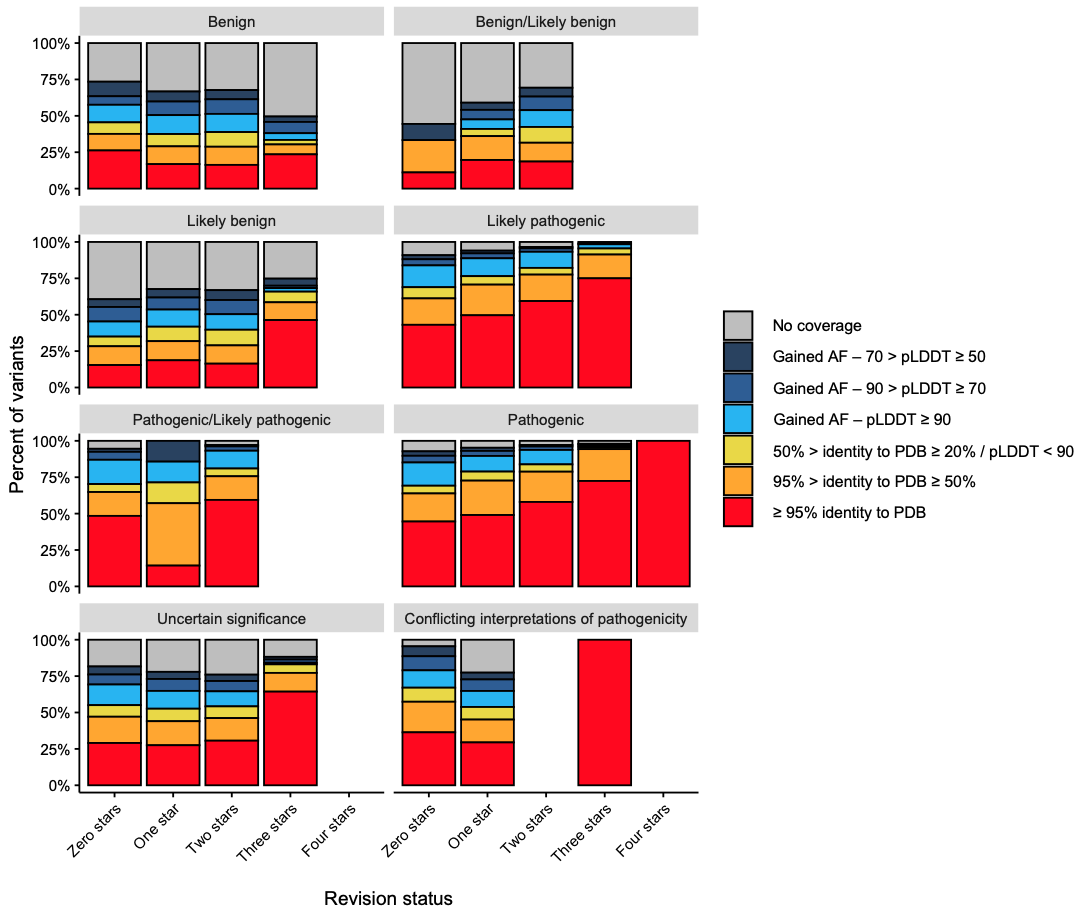

**Supplementary Figure 1 –** Structural coverage of Clinvar mtuations (y-axis) depending on their pathogenicity (different panels) and their review status (x-axis)

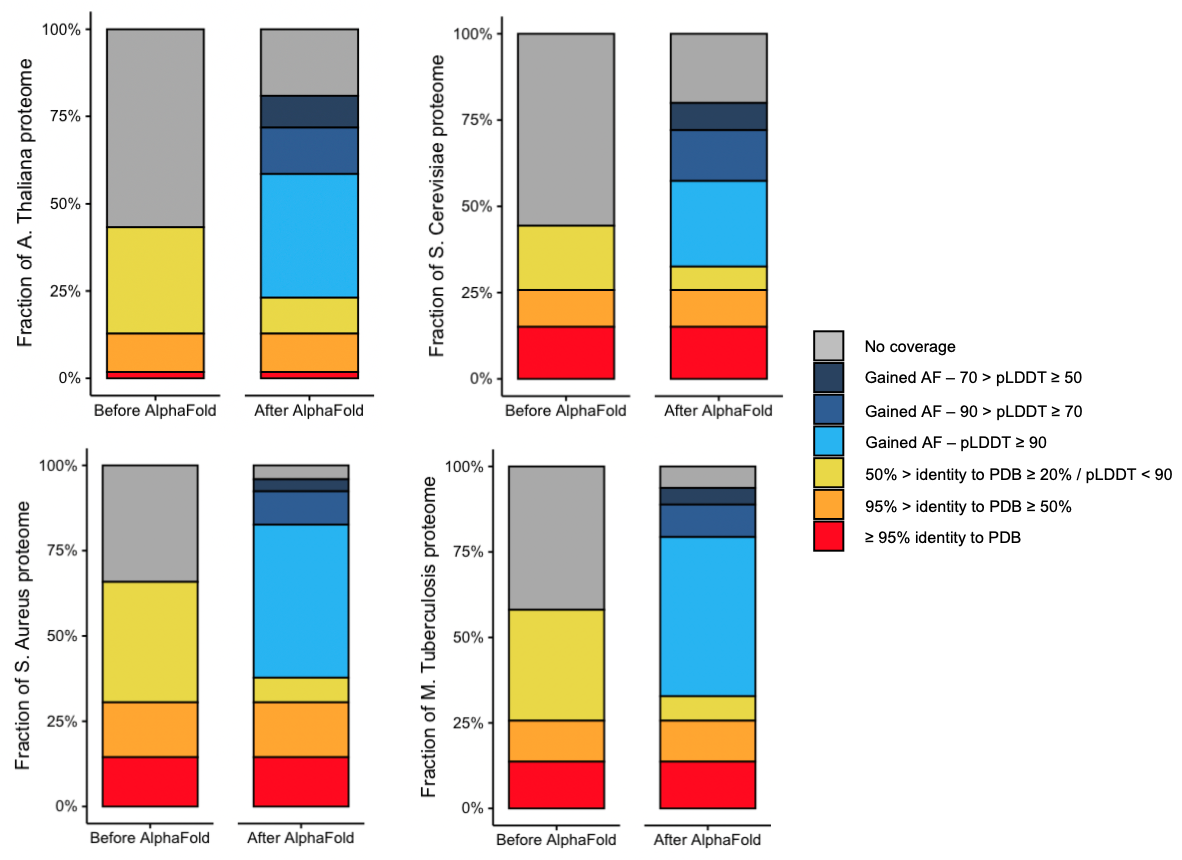

**Supplementary Figure 2 –** Structural coverage (y-axis) of the proteome of the four different organisms before (left) and after (right) including the AlphaFold models
